# Accounting for the influence of dissimilarity gradients on community uniqueness

**DOI:** 10.1101/2025.09.28.679068

**Authors:** Daniel Hernández-Carrasco, Anthony J. Gillis, Hao Ran Lai, Tadeu Siqueira, Jonathan D. Tonkin

**Affiliations:** School of Biological Sciences, University of Canterbury, Christchurch 8140, New Zealand; South East Asian Rainforest Research Partnership (SEARRP), Kota Kinabalu, Malaysia; Institute of Biosciences, São Paulo State University (UNESP), Rio Claro, SP, Brazil; Te Pūnaha Matatini Centre of Research Excellence, University of Canterbury, Christchurch 8140, New Zealand

## Abstract

Compositional uniqueness has become increasingly relevant for understanding how local communities contribute to regional biodiversity. The most widely used metric is the Local Contribution to Beta Diversity (LCBD), which is typically regressed against environmental predictors. However, LCBD can vary either because of environmental processes that affect the overall variance in community composition, or because communities change directionally along environmental gradients. The latter implies that LCBD–environment relationships can strongly depend on how the environment is sampled. To address this issue, we introduce Generalised Dissimilarity Uniqueness Models (GDUM), a framework that embeds effects on community uniqueness within pairwise dissimilarity modelling. GDUMs are consistent with conventional uniqueness models, while explicitly accounting for directional changes in composition. This distinction disentangles directional and non-directional drivers of beta diversity, such as environmental filtering versus stochastic processes. By improving interpretability and generalizability, GDUM is a useful tool for understanding beta diversity patterns and projecting biodiversity responses.

## 1 Introduction

Beta diversity, the variation in community composition across space and time, has become a cornerstone of community ecology. It has helped uncover patterns in community structure (Anderson et al., 2011; Villarino et al., 2022; Zellweger et al., 2017), disentangle mechanisms driving community assembly (Chase, 2010; Dobrovolski et al., 2012; Jacobi & Siqueira, 2023; Soininen et al., 2018), and detect the footprint of human impact on biodiversity (Gutiérrez-Cánovas et al., 2013; Socolar et al., 2016; Su et al., 2021). These broad applications have sparked increasing interest in improving statistical tools (Dias et al., 2022; Mokany et al., 2022; White et al., 2024). Conceptually, beta diversity encompasses two complementary interpretations (Anderson et al., 2011): directional community change, where communities change predictably along environmental or spatial gradients, and non-directional variation, where the focus is on the overall dispersion of community composition (Fig. 1A). While recent methodological advances have focused on capturing directional changes, metrics related to overall community variance and the contribution of individual sites have been analysed separately. This separation hinders both inference and prediction about biodiversity change.

**Fig. 1:**
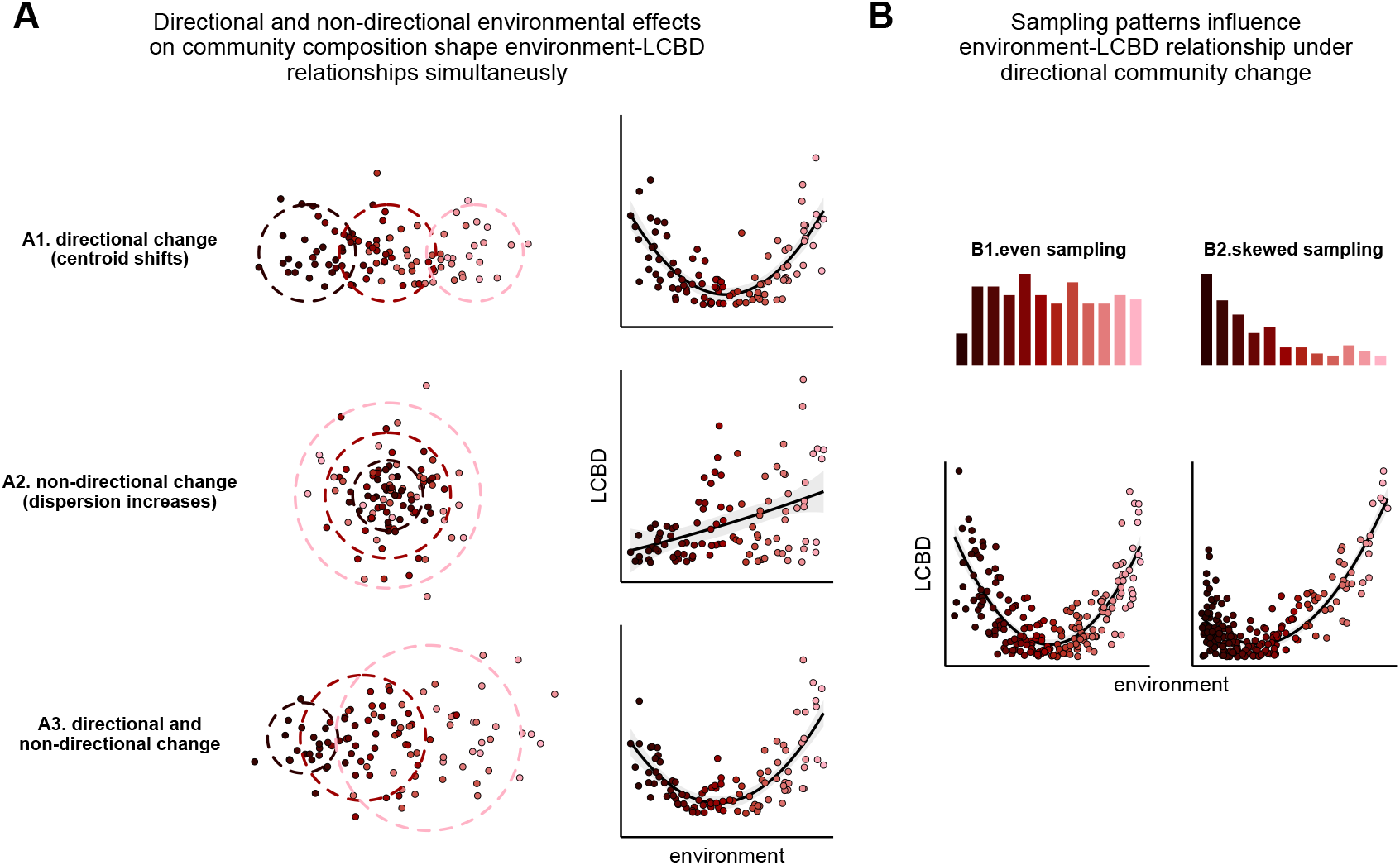
LCBD-environment relationships depend on directional and non-directional environmental effects on community composition, and how environmental gradients have been sampled. Colour gradient represents a hypothetical environmental variable. (A) Example of three patterns of variation in community composition driven by directional and/or non-directional environmental effects (left), and resulting environment-LCBD relationships (right). (B) Effect of sampling patterns (top) on environment-LCBD relationship (bottom) under directional community change.

Understanding how individual sites contribute to spatial biodiversity patterns and overall beta diversity can provide insight into local processes that shape the site’s relative uniqueness. The uniqueness of a community within a region can be quantified as its average dissimilarity to other communities, its distance to the centroid in an ordination space, or more commonly, as the community’s Local Contribution to Beta Diversity (LCBD) (Anderson et al., 2006; Legendre & De Cáceres, 2013). LCBD indices have been used as a tool for identifying high priority conservation sites across biomes, habitats, and taxa (Dubois et al., 2020; Fernández-Calero et al., 2024; Heino & Grönroos, 2017; Hill et al., 2021; Liu et al., 2024; Rodríguez-Lozano et al., 2023; Yan et al., 2021), and for detecting the impact of localised stressors (Landeiro et al., 2018; Schneck et al., 2022; Shi et al., 2024). Unravelling the causes of compositional uniqueness can therefore aid conservation planning and shed light on mechanisms governing biodiversity change. For instance, high LCBD values have been linked to ecological drift, as well as spatial processes including habitat isolation and dispersal limitation (Fernández-Calero et al., 2024; Li et al., 2020; Lozada et al., 2023). Similarly, the degree of invasion by non-native species can be an important driver of variability in compositional uniqueness (Bando et al., 2023). These processes alone can explain a proportion of the variability observed in LCBD, but current tools for analysing community uniqueness do not yet explicitly account for an important set of mechanisms: those generating directional changes in community composition.

While LCBD may appear as an individual site-level metric, it is based on *pairwise* dissimilarities between the focal and other sites. Consequently, environmental gradients such as elevation, latitude, or nutrient concentration (Qian & Ricklefs, 2012; Wang et al., 2012) that produce directional changes in community composition (i.e., turnover *sensu* Anderson et al., 2011), can influence LCBD by shaping the underlying dissimilarity matrix. Environmental effects on LCBD may therefore reflect the availability of those environmental conditions in the landscape or their sampling frequency, resulting in arbitrarily complex LCBD–environment relationships (Fig. 1B) (Tsang et al., 2023). For instance, continuous community change across an evenly sampled environmental gradient can produce a U-shaped LCBD–environment relationship because communities at the extremes of the gradient would be relatively rare, such as at either end of an elevational gradient (Wang et al., 2020; Yang et al., 2022). However, if more samples are taken at the ends of the gradient, the observed pattern could be reversed. The fact that such seemingly contrasting relationships can arise along the same gradient suggests the importance of disentangling directional vs. non-directional environmental effects on compositional change.

Here, we present Generalised Dissimilarity Uniqueness Models (GDUM), a multi-level regression framework that extends previous community dissimilarity models by enabling direct estimation of uniqueness effects, which we show are mathematically equivalent to conventional LCBD approaches. Because GDUM is a type of dissimilarity model, it also accommodates pairwise predictors to capture directional community change along environmental gradients (Mokany et al., 2022). Thus, GDUM allows simultaneous modelling of the drivers of dissimilarity gradients and site-level uniqueness, thereby improving estimates of both. We use simulated data to test the ability of the proposed model to disentangle dissimilarity gradients and direct effects on compositional uniqueness, and then demonstrate the applicability of our approach with two empirical case studies.

## 2 Generalised Dissimilarity Uniqueness Models

### 2.1 Model overview

We define a model for the element *y*_*ij*_ of a square matrix **Y**, representing the dissimilarity between site *i* and *j* (with *i, j* ∈ {1, 2, · · ·, *n*} and *i* > *j*) as follows:

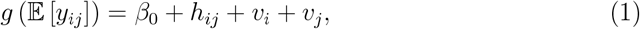

where *g*(·) denotes a link function mapping the mean of an assumed distribution (often a binomial or beta distribution for dissimilarity matrices bounded between 0 and 1) to a linear predictor. In Equation (1), *β*_0_ is a global intercept. As indicated by their indices, *h*_*ij*_ is a pairwise sub-model, whereas *v*_*i*_ and *v*_*j*_ are site-level components as row and column sub-models.

For the pairwise sub-model, let *x*_*ik*_ be the value of the *k*-th environmental predictor of site *i* from a matrix of predictors **X**. Following Ferrier et al. (2007), we define *h*_*ij*_ as a function of the distance between predictors after applying a non-linear transformation

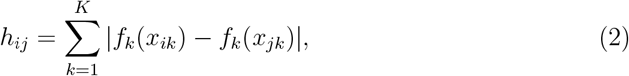

where *f* (·) are usually defined as monotonic splines to capture non-linear community change (Appendix S1; see Mokany et al., 2022 for a detailed discussion of the approach). For the case where *f* (·) is the identity function, *h*_*ij*_ can be simply written as *h*_*ij*_ = |**x**_*i*_ − **x**_*j*_|***β***, where ***β*** is a column vector of coefficients.

For the row and column sub-models (site component), let **w**_*i*_ be the *i*-th row of a second matrix of environmental predictors **W, *λ*** be a column vector of coefficients, and *z*_*i*_, *z*_*j*_ represent site-level random effects:

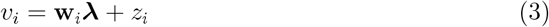

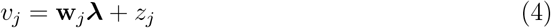

We note that ***λ*** appears in both Equations 3 and 4, representing the same effect of the same environmental predictor on both sites of the pair. Likewise, *z*_*i*_ and *z*_*j*_ represent two elements from the same vector of random effects ***z***. The row and column sub-models determine the average dissimilarity of each site to all other sites, and therefore intuitively relates to their uniqueness. A formal interpretation is provided in the following sub-section.

### 2.2 Link to a conventional site uniqueness model

The Local Contribution to Beta Diversity (LCBD) is computed by double-centring the pairwise dissimilarity matrix **Y**. This yields a matrix **G** whose diagonal entries *g*_*ii*_ are site-specific sums of squares (SS_*i*_) interpreted as uniqueness scores (Legendre & De Cáceres, 2013). The double-centring procedure corresponds to the first step in a Principal Coordinates Analysis (PCoA) (Gower, 1966), where SS_*i*_ = *g*_*ii*_ represents the multivariate distance of each site to the centroid. LCBD scores are obtained after scaling SS_*i*_:

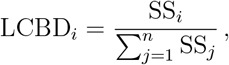

but we consider unscaled SS_*i*_ values in the following for simplicity. Site uniqueness is commonly modelled as a function of site-level covariates (Hill et al., 2021; Montrás-Janer et al., 2024; Ochieng et al., 2024) with the linear predictor

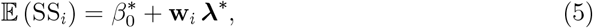

where **w**_*i*_ is the vector of environmental predictors at site *i*, ***λ***^∗^ are their associated model coefficients and 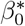 is the intercept. We show that a simple mathematical relationship exists between the coefficients in the site uniqueness model (Equation 5) and those in the site-level component of GDUMs (see full proof in Appendix S2):

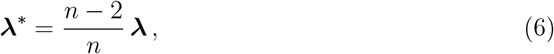

which implies that site-level effects in the *v*_*i*_ and *v*_*j*_ sub-models are equivalent to those in a conventional site uniqueness model. This mathematical relationship provides a clear interpretation of site-level model coefficients in GDUMs, linking them directly to environmental effects on community uniqueness. Moreover, site-level random effects (*z*_*i*_, *z*_*j*_) can be related to the error term in the site uniqueness model (Appendix S2).

### 2.3 Estimation

In the accompanying R package `gdmmTMB`, model parameters are estimated using maximum likelihood. We implement the model in C++ using the `Template Model Builder` (TMB) framework, which provides efficient marginal likelihood evaluation (Kristensen et al., 2016). Optimization is carried out in R using the `nlminb` algorithm from the `stats` package. We consider two alternative strategies for parameter estimation, both used in previous versions of dissimilarity models to account for non-independencies in dyadic (pairwise) data (Dias et al., 2022; Woolley et al., 2017):

#### 1. Full hierarchical model

Site-level random effects are included explicitly, capturing site-to-site heterogeneity. Classic gradient-based standard errors for model parameters are computed via `sdreport` in TMB. For derived quantities such as dissimilarity and uniqueness, we implement parametric bootstrapping to obtain robust credible intervals. In brief, parametric bootstrapping consists in drawing *S* samples from the multivariate normal distribution defined by the estimated parameters 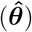 and their covariance matrix 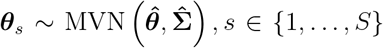 Expected dissimilarity and uniqueness are then recomputed for each draw, and the resulting distribution is used to construct credible intervals.

The full hierarchical model definition is the default strategy used in `gdmmTMB` to estimate model parameters. However, constraints in model coefficients—such as those required to fit monotonic splines—can complicate the calculation of standard errors from the Hessian, in which case we recommend the Bayesian bootstrapping approach (see below).

#### 2. Bayesian bootstrapping

Random weights are assigned to each observation (site) *i* and used in the computation of the log-likelihood. We repeat this process by fitting the model with different weight combinations and analyse the empirical distribution of parameters to quantify uncertainty in both model parameters and derived quantities. This procedure is analogous to common bootstrapping with replacement, where the model is fitted with different permutations of the original data. We note that, while Bayesian bootstrapping avoids the need to estimate the variance–covariance matrix, it requires refitting the model multiple times, which can be computationally intensive.

### 2.4 Dissimilarity metrics

Because the response variable in GDUM is a dissimilarity index, special attention must be paid to the link function and error distribution to ensure that the mathematical properties of the selected index are appropriately accounted for. For instance, many dissimilarity indices commonly used in community ecology represent proportions of shared species or individuals and are therefore bounded between zero and one. In those cases, a beta distribution is a natural choice (Dias et al., 2021). An alternative approach is to recognise that some dissimilarity indices can be expressed as ratios of counts, which makes them compatible with a binomial likelihood (Woolley et al., 2017). For example, the Sørensen–Dice dissimilarity represents the proportion of unshared species relative to the total number of species across a pair of sites:

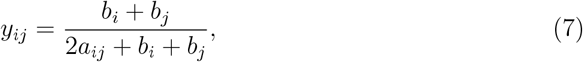

where *a*_*ij*_ is the number of shared species between sites *i* and *j*, and *b*_*i*_ and *b*_*j*_ are the numbers of species unique to each site. The expected Sørensen–Dice dissimilarity can therefore be modelled as the probability *π* = 𝔼 (*y*_*ij*_) that a randomly chosen species from the combined pool is not shared between the two sites:

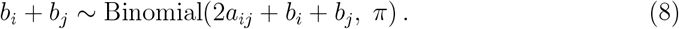

For the purposes of the case studies presented here, we use the Sørensen–Dice dissimilarity index and model it using a binomial distribution and the canonical logit link function. However, different options for dissimilarity indices, error distributions and link functions can be accommodated within the GDUM framework—many of them supported in `gdmmTMB`.

## 3 Simulation study

We compared the ability of GDUM to recover direct effects on uniqueness to that of conventional uniqueness models with a simulation study. Data were generated from a “true” model (Equation 1) with known parameter values for *λ* and *β*, and a range of realistic sample sizes. For each combination of parameter values and sample sizes, we generated 1000 replicate datasets. We considered two field-sampling scenarios regarding the sampled distribution of the predictor *x*_*i*_: (1) normally distributed values or (2) slightly skewed distribution, where small values were less frequently sampled (Fig. 2A-B). Each simulated dataset was analysed using three competing models: a GDUM with site-level random effects (Equation 1), a GDUM estimated with Bayesian Bootstrapping, and a conventional model of SS_*i*_ (Equation 5). We scaled the true parameters from the latter using Equation (6) to enable the comparison between different models. For each fitted model, we quantified bias (mean difference between estimated and true values), root mean square error (RMSE), and evaluated inferential performance by estimating coverage probabilities of 95% confidence or credible intervals. Site-level and pairwise-level error terms were drawn from Gaussian distributions with standard deviation *σ* = 0.5.

**Fig. 2:**
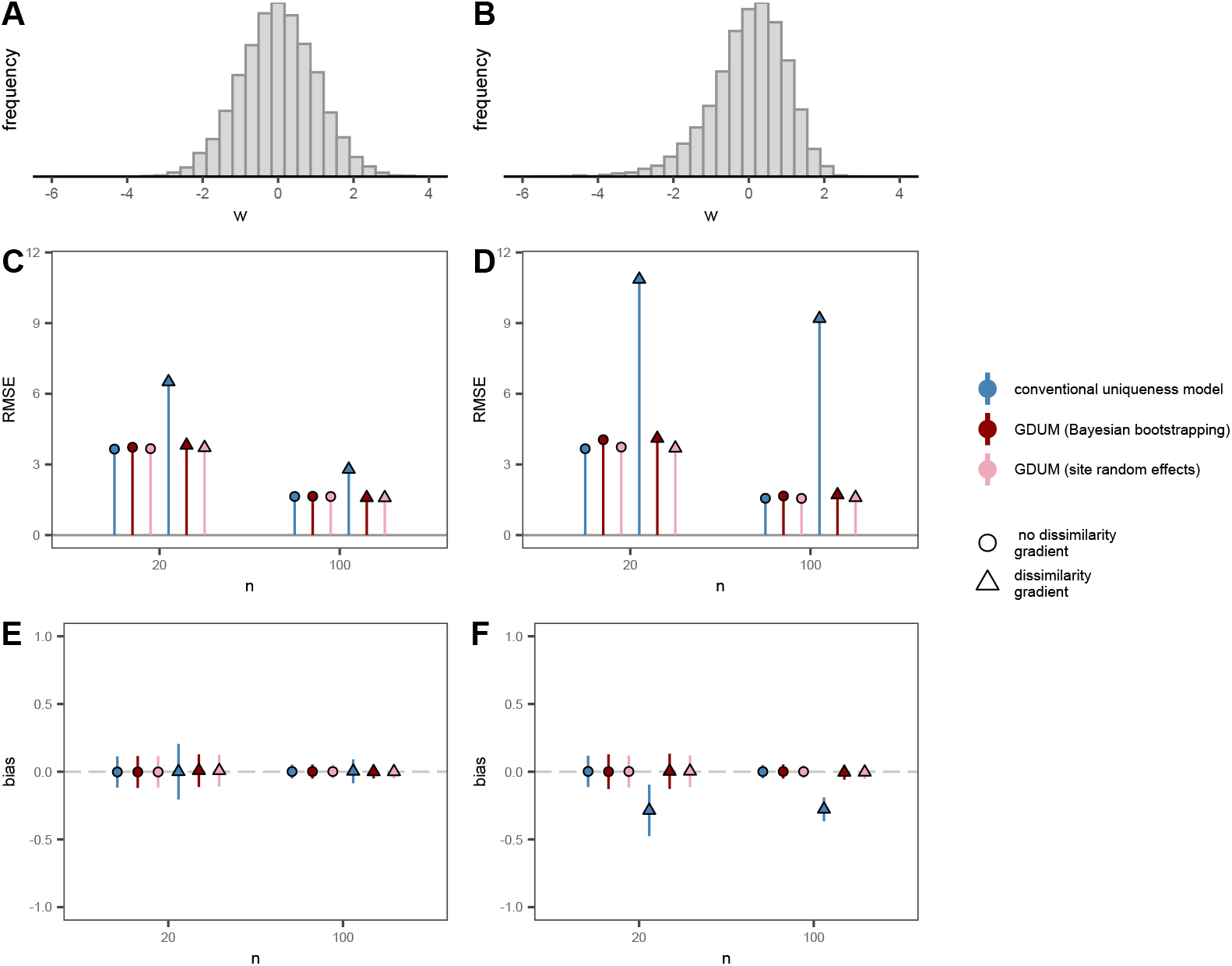
Comparison of model performance in our simulation study. Each model contained *n* values of a predictor *w* sampled from a symmetric, Gaussian distribution (A, C, D) or a slightly skewed distribution (B, D, F). We considered cases where (1) *w* only had a direct effect on uniqueness (*no dissimilarity gradient* ; *β* = 0) and (2) *w* was also associated with a dissimilarity gradient (*dissimilarity gradient β* = 1). Panels (C) and (D) show model error as the Root Mean Square Error (RMSE) between the recovered regression parameter *λ* and the true simulated parameter, for models fitted with the symmetric and skewed predictor, respectively. Panels (E) and (F) show the bias of recovered *λ* coefficients as the average deviation from the true parameter. Error bars indicate one standard deviation from the average.

The error associated with the estimated uniqueness effect (*λ*) was similar for all three models in scenarios where dissimilarity gradients were not included (*β* = 0). However, RMSE was consistently higher for the conventional site uniqueness model when dissimilarity gradients were present (Fig. 2), particularly when the predictor had a skewed distribution. Moreover, the slight skewness in the predictor variable introduced a systematic underestimation of *λ* values in the conventional site uniqueness model, while estimates from GDUMs were largely unaffected. GDUMs performed similarly in scenarios with and without dissimilarity gradients. Confidence intervals aligned with the expected 95% true parameter coverage (for *α* = 0.05) for GDUMs with site-random effects, except for small sample sizes (*n* = 20), which showed slightly lower coverage rates (Fig. S2, S3). Overall, the full hierarchical implementation of GDUM showed the best performance in terms of bias, RMSE, and parameter coverage across all scenarios.

## 4 Case studies

We demonstrate the applicability of GDUM with two case studies. First, we show how strong directional patterns of community change along an environmental gradient can drive the observed patterns of community uniqueness, and how GDUMs can account for such effects. Second, we showcase the use of our approach for studies with multiple predictors.

### 4.1 A cautionary tale: Microbial communities along a pH gradient

To demonstrate the potential influence of directional community change on environment– LCBD relationships, our first example centres on community composition along a relatively steep environmental gradient. We used a dataset of microbial communities available as part of the gllvm R package (Niku et al., 2025), which contains a known pattern of change in species composition from low to high pH levels (Niku et al., 2017). The dataset was originally published in Nissinen et al. (2012), and contains presence– absence information for 985 species across 56 sites. Although the dataset contains multiple environmental variables, we focused on pH as the main driver of community change.

As is standard in the literature (Da Silva et al., 2018; García-Navas et al., 2022; Valente-Neto et al., 2020), we first computed LCBD scores from the dissimilarity matrix and regressed them against pH and the quadratic pH term. To represent how compositional changes along the pH gradient influence the pH–LCBD relationship, we repeated the process using the residual dissimilarity after accounting for directional community change within the *h*_*ij*_ component of a dissimilarity model. We performed a Non-Metric Multidimensional Scaling (NMDS) with two dimensions on (1) the raw dissimilarity matrix and (2) the residual dissimilarity to visualise this effect. For comparison, we also fitted a full GDUM with both an environmental dissimilarity gradient and a direct environmental effect on uniqueness.

The conventional uniqueness model (Fig. 3C) showed a U-shaped relationship between LCBD and pH, driven by the strong, directional change in community composition along the pH gradient (Fig. 3A). By contrast, accounting for such directional change revealed that the non-directional effect of pH on uniqueness had the opposite shape, with intermediate pH being associated with higher community uniqueness (Fig. 3D). This pattern was also reflected in the ordination of residual dissimilarities (Fig. 3C), where samples at both ends of the pH gradient appeared closer to the centroid. The full GDUM approach successfully captured both the expected U-shaped relationship driven by directional community change, and the humped-shaped relationship driven by the increased variability in community composition at intermediate pH levels (Fig. 4C). Combined, both components provided an expected pH-LCBD relationship that adjusted remarkably well to the observed pattern (Fig. 4A). Our approach therefore overcomes a crucial limitation of uniqueness models (Tsang et al., 2023), namely, distinguishing between direct effects on uniqueness (non-directional effects) and patterns caused by a combination of directional community change and arbitrary sampling schemes.

**Fig. 3:**
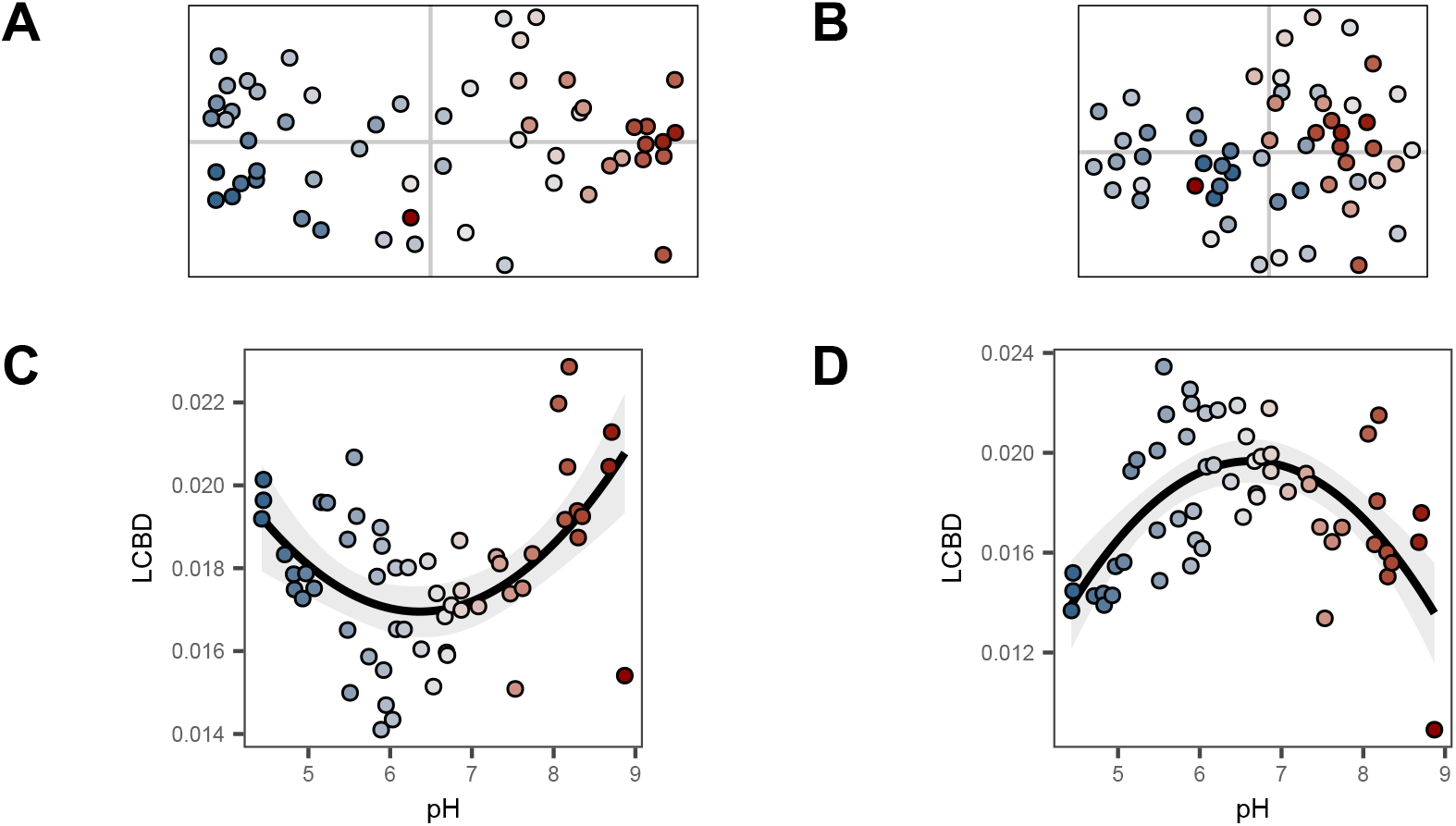
Empirical pattern of compositional uniqueness along a steep environmental gradient (pH) is driven by directional community change. (A) Variability in microbial community composition represented by a Non-Metric Multidimensional Scaling (NMDS). (B) Variability in community composition after accounting for directional community change along the pH gradient. Residual dissimilarities were obtained after fitting a dissimilarity model with a dissimilarity gradient along (*h*_*ij*_). (C) Empirical LCBD-pH relationship. The shape of a quadratic regression is displayed with associated standard error. (D) LCBD-pH relationship after accounting for directional change in community composition.

**Fig. 4:**
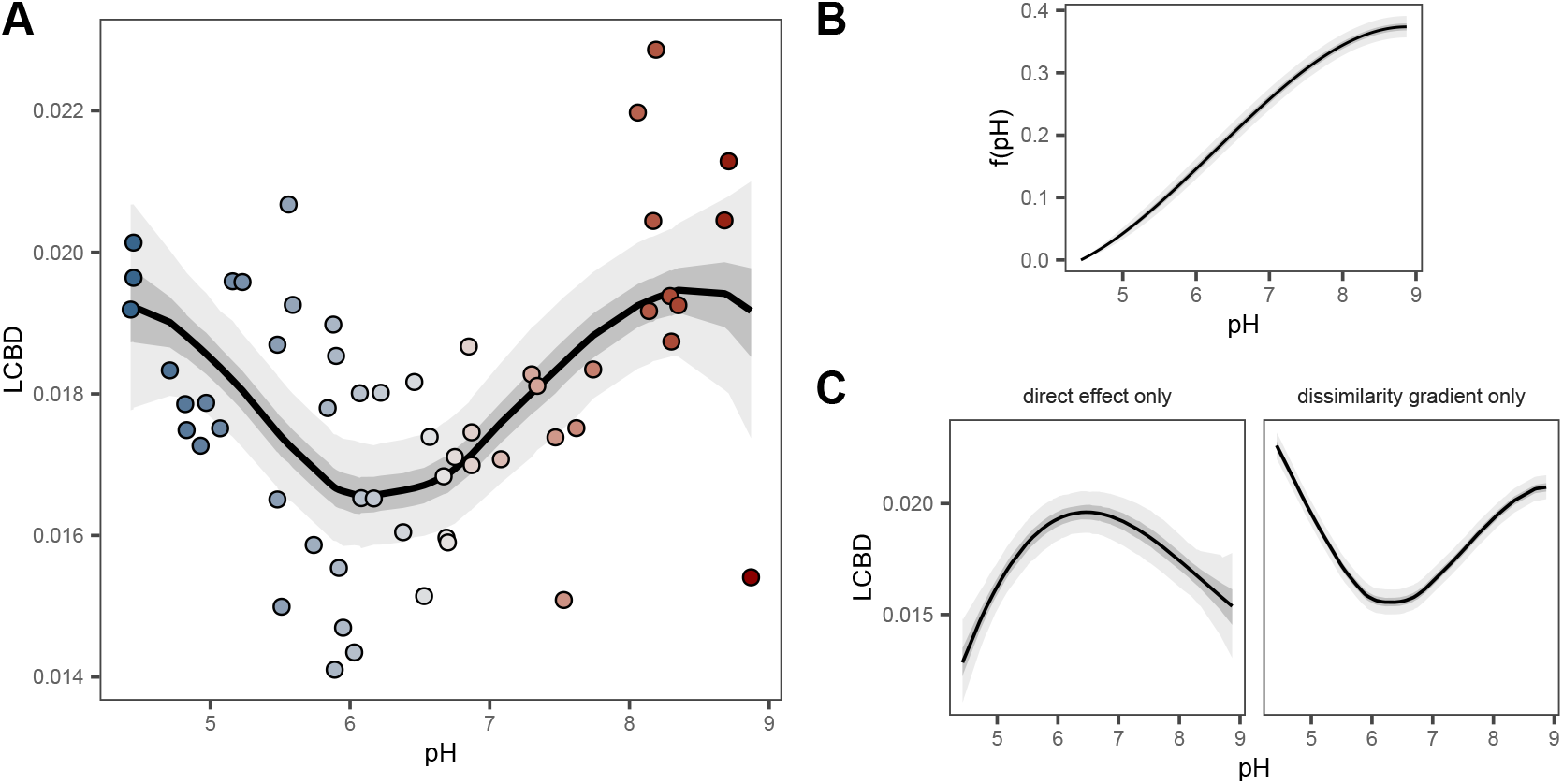
Contribution of pairwise and site-level environmental effects to the expected LCBD. (A) Relationship between LCBD and pH obtained from a GDUM with both a dissimilarity gradient and direct effects on uniqueness. Circles represent *empirical* values computed using the LCBD.comp function in the adespatial R package (Dray et al., 2025). Expected LCBD values from GDUM were derived from the expected dissimilarity matrix. (B) Fitted dissimilarity gradient showing the directional community change with pH. (C) Partial effect of pH on LCBD when included only as a site-level variable (*v*_*i*_, *v*_*j*_; left) or as a dissimilarity gradient (*h*_*ij*_; right).

The contribution of the fitted dissimilarity gradient (Fig. 4B) to the observed uniqueness pattern depends not only on the estimated coefficients in *h*_*ij*_, but also on the distribution of the environmental variable (Fig. S4). This property of the GDUM framework provides the necessary flexibility to project community uniqueness under counterfactual sampling schemes. For instance, projections given new environmental data can represent the availability of different environmental conditions across the whole landscape (rather than the sampled locations only), alternative management scenarios, or potential environmental changes.

### 4.2 Usage example: Plant community change along environmental gradients

Our second example demonstrates the applicability of the GDUM approach to broader studies. We used a dataset comprising plant community composition from South-West Australia available as part of the `gdm R` package (Fitzpatrick et al., 2025). The dataset contains information on species presence-absence, as well as local and bioclimatic environmental predictors compiled from multiple sources. We limited the scope of the cae study to four bioclimatic predictors: BIO5 (maximum temperature of warmest month), BIO6 (minimum temperature of the coldest month), BIO15 (precipitation seasonality) and BIO19 (precipitation of the coldest quarter).

We included the four predictors in both the pairwise (as **X** in *h*_*ij*_) and site level components (as **W** in *v*_*i*_, *v*_*j*_), and estimated their effects using Bayesian bootstrapping. We decomposed variables in the pairwise component into 3 I-spline basis of degree 2 (*f*_*k*_(·)) to account for non-linear community change (Ferrier et al., 2007) except for spatial distance (*d*_*ij*_), which we included directly as a pairwise variable:

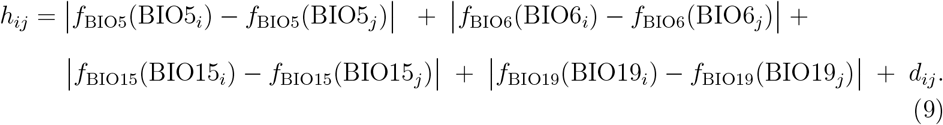

As is common in dissimilarity models (Ferrier et al., 2007; White et al., 2024), we defined *f*_*k*_(·) as monotonic I-splines by constraining associated coefficients to the positive domain. This constraint ensures parameter identifiability and improves the interpretation of the pairwise component. We assumed linear site-level effects in the row and column sub-models (i.e., linear effects on site uniqueness):

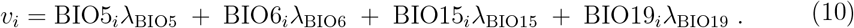

Model results showed strong directional community change along variables BIO19 and BIO5 (Fig. 5A). Specifically, the representation of the dissimilarity gradients revealed higher rates of community change at low BIO19 values, whereas community change along BIO5 was relatively linear. Although directional community change along BIO15 was relatively small, it had a significant site-level effect on uniqueness (Fig. 5B). Overall, the inclusion of dissimilarity gradients provided a better fit for both pairwise dissimilarities and site uniqueness (Fig. 5C-D, Fig. S5C-F), and had a strong influence on recovered site-level effects (Fig. S5A-B).

**Fig. 5:**
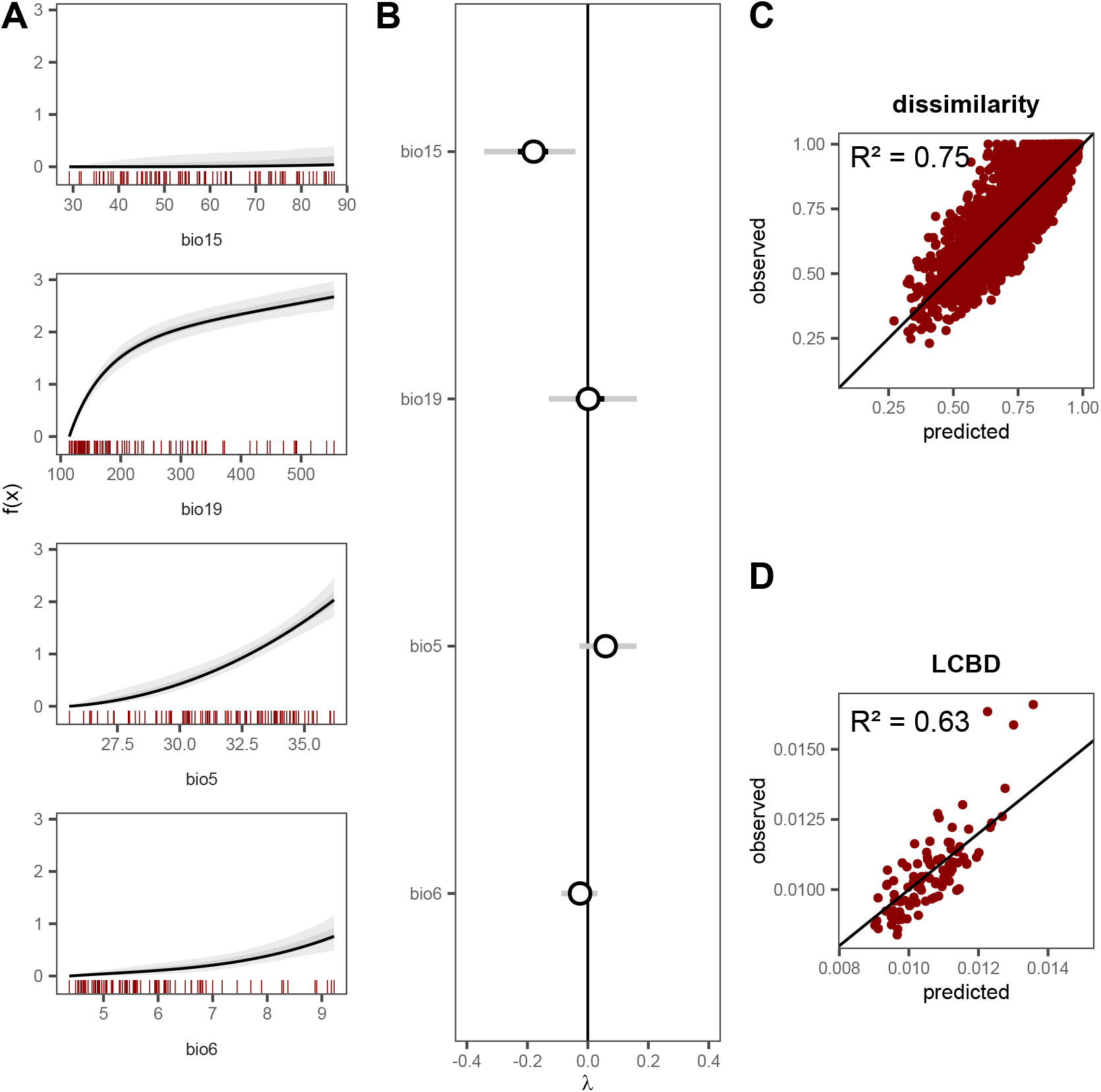
Dissimilarity gradients along bioclimatic variables and direct effects on uniqueness. (A) Dissimilarity gradients representing directional community change. *f* (*x*) represents the value of environmental predictors after applying the fitted non-linear transformation with monotonic I-splines. Red vertical lines indicate the empirical distribution of each variable in the data. (B) Estimated partial effects of bioclimatic variables on community uniqueness. Error bars indicate 95% and 50% credible intervals. Panels (C) and (D) show the relationship between predicted and observed dissimilarities and LCBD values, respectively. *Observed* LCBD values represent empirical scores computed with the LCBD.comp function in the adespatial R package (Dray et al., 2025).

## 5 Discussion

Developing robust frameworks for modelling beta diversity is crucial for understanding and predicting biodiversity change. Here, we introduce Generalised Dissimilarity Uniqueness Models (GDUM), an extension of previous dissimilarity models that enables the inclusion of direct effects on community uniqueness. In doing so, GDUMs address severe limitations of current approaches to uniqueness, and unify two facets of beta diversity: directional community change along gradients and non-directional change (Anderson et al., 2011).

If not accounted for, dissimilarity gradients can lead to biased site-level effects (i.e., direct effects on uniqueness) under directional community change. This was reflected in our simulations, where even mildly skewed environmental predictors led to systematic bias in parameter estimation. We note that non-linear community changes would lead to qualitatively similar results even if predictors were not skewed. Transforming the variables prior to model fitting would therefore not solve this issue in realistic scenarios (D’Amario et al., 2019; Xiong et al., 2020). Thus, variables associated with directional changes in community composition can lead to spurious uniqueness–environment relationships in conventional uniqueness models that depend on their sampled distribution (Tsang et al., 2023). For instance, U-shaped relationships between LCBD and environmental variables associated with strong species turnover, such as elevation, have been repeatedly reported (Ochieng et al., 2024; Ren et al., 2024; Vad et al., 2017). Similarly, linear relationships are often found when the distribution of the variable of interest is strongly skewed (García-Navas et al., 2022; Valente-Neto et al., 2020). These relationships represent the real correlations in the data, but unless the role of directional community change is explicitly accounted for, it remains unclear whether they reflect true ecological processes or stem from arbitrary sampling schemes. Moreover, because environment-uniqueness relationships are contingent on the sampled distribution of environmental predictors, they may shift under novel conditions, as environments that were relatively rare can become more common and vice-versa (Vasconcelos et al., 2018; Williams & Jackson, 2007). Our approach offers a solution to this widespread issue by explicitly incorporating gradients of directional community change—which allows recalibrating uniqueness estimates as new data arrives. As a result, GDUM provides a more robust framework for projecting compositional uniqueness under environmental change scenarios, and are better suited for scenario testing of management or conservation interventions.

Alternative methods have been proposed to address the influence of directional community change on recovered environment-uniqueness effects, most notably: (1) incorporating ‘environmental uniqueness’ as a measure of environmental rarity (Snåre et al., 2024), and (2) simulating homogeneous sampling scenarios after fitting a dissimilarity model (Tsang et al., 2023). However, both strategies pose important limitations. Environmental uniqueness can be informative, but it is typically calculated across all predictors simultaneously, which may obscure the role of individual environmental drivers. This approach also requires the assumption of linear community change, and can lead to multicollinearity when both the raw predictors and their rarity are included as they would be strongly correlated under many sampling scenarios. Simulating homogeneous sampling offers a potential improvement, but the influence of dissimilarity gradients is only fully removed if community change is strictly linear, and it is unclear how direct effects on ecological uniqueness should be included.

Unlike conventional approaches, the response variable in GDUM is a pairwise dissimilarity, so accounting for directional community change is straightforward. In our case studies, we included environmental gradients as absolute differences after a non-linear transformation (Ferrier et al., 2007; Mokany et al., 2022). Non-linear transformations of pairwise predictors, such as splines, can capture differences in the rate of community change along environmental gradients, offering great flexibility. In addition to environmental variables, the pairwise component of GDUMs can capture spatio-temporal dependencies in community composition. For instance, incorporating a distance decay of similarity can capture unexplained spatial autocorrelation (Dias et al., 2021; Soininen et al., 2007) as shown in our second case study.

In addition to accounting for dissimilarity gradients, GDUMs incorporate direct effects of local processes on compositional uniqueness with site-level predictors. We have shown that these effects are mathematically equivalent to those of predictors in conventional uniqueness models. Good candidates for site-level predictors are variables that could affect community uniqueness through processes beyond directional changes, by increasing or decreasing the average dissimilarity between a given site and all other sites. That is, by increasing community variance (beta diversity) in a non-directional manner (*sensu* Anderson et al., 2011) (Fig 1A). For instance, spatial isolation can lead to more unique assemblages due to the small influence of surrounding communities compared to more connected sites. Highly isolated communities are not necessarily more similar to other isolated communities, but rather present distinct assemblages compared to all other habitats (higher variance), ultimately leading to higher uniqueness (Fernández-Calero et al., 2024; Lozada et al., 2023). Other examples of variables with potential effects on compositional uniqueness include habitat size, which can modulate the effect of ecological drift (Green et al., 2022; Lozada et al., 2023), herbivory, through the lower occurrence rate of rare taxa (Villar et al., 2020), and productivity, through an increased variability in community composition (Chase, 2010).

## 6 Conclusions

Quantifying the uniqueness of ecological communities–a key facet of beta diversity–and correctly inferring the role of environmental drivers is central to both basic and applied ecology, particularly for conservation planning. However, conventional modelling approaches can obscure the underlying causes of uniqueness by inflating, downplaying, or confounding relationships with environmental predictors. By disentangling directional from non-directional effects, GDUMs overcome this limitation and unlock new opportunities to investigate how specific drivers shape beta diversity patterns. Moreover, GDUMs offer the necessary flexibility to project uniqueness to non-sampled locations, future environments, and different management strategies, making them a powerful tool for applied studies.

## 7 Acknowledgements

This research was funded by a Rutherford Discovery Fellowship administered by the Royal Society Te Apārangi (RDF-18-UOC-007) awarded to JDT. JDT is also supported by Te Pūnaha Matatini, a Centre of Research Excellence funded by the Tertiary Education Commission, New Zealand. HRL is supported by the Marsden Fund managed by the Royal Society Te Apārangi (grant MFP-UOC2102) and the Bioprotection Aotearoa Centre of Research Excellence. TS is supported by the Conselho Nacional de Desenvolvimento Científico e Tecnológico - CNPq/Brazil (Productivity Grant, 309496/2021-7). DHC receives funding from the University of Canterbury Doctoral Scholarship.

## Author contribution

DHC, HRL and AJG conceived the ideas and designed methodology. DHC led the data analysis and the writing of the manuscript. JDT obtained funding. All authors contributed critically to the drafts and gave final approval for publication.

## Data availability

Data used in the two case studies are publicly available as part of existing software archived in The Comprehensive R Archive Network (cran.r-project.org).

Soil microbial data are available as part of the gllvm R package (10.32614/CRAN.package.gllvm) and plant community data are available as part of the gdm package (10.32614/CRAN.package.gdm). Code used to reproduce the results and figures in this manuscript is available on GitHub (github.com/dhercar/GDUM) and has been archived in Zenodo (doi.org/10.5281/zenodo.17128110). An R package for fitting Generalised Dissimilarity Uniqueness Models is available on GitHub (github.com/dhercar/gdmmTMB)

