## Supporting Information for "Accounting for the influence of dissimilarity gradients on community uniqueness"

#### Appendix S1: Visualising dissimilarity gradients

Patterns of directional community change modelled within the pairwise component of a GDUM ($h_{ij}$) can be visualised by plotting each raw predictor $x$ against their (fitted) transformed value $f_x(\cdot)$ (Fig. S1). The contribution of each dissimilarity gradient to the linear predictor is the absolute difference between transformed predictors, which makes the shape of the resulting function suitable for visualising the rate of community change at different values of $x$. In addition, the magnitude of $f_x(x)$ for different predictors can be interpreted as a proxy of their relative importance.

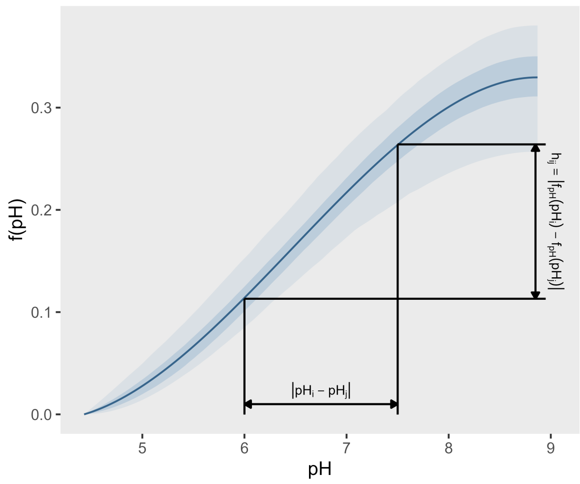


Fig. S1: Non-linear dissimilarity gradient fitted with monotonic I-splines. Coloured areas represent 95% and 50% credible intervals. In this example, the contribution of the pairwise component $h_{ij}$ to the linear predictor is the absolute difference of pH values after the non-linear transformation $f(\cdot)$ has been applied.

### Appendix S2: Mathematical relationship between GDUM and the conventional uniqueness model

#### 2.1 Model formulation and notation

##### 2.1.1 Pairwise dissimilarity model

Consider a dataset comprising $n$ sites with pairwise dissimilarity matrix $\mathbf{Y}$ and environmental predictor matrix $\mathbf{W}$, where $\mathbf{w}_i$ represents the row vector of predictors for site $i$. Let $\boldsymbol{\lambda}$ denote the column vector of regression coefficients and $\beta_0$ the intercept term. The pairwise dissimilarity model is defined for each element $y_{ij}$ of $\mathbf{Y}$ as:

$$\mathbb{E}\left(y_{ij}\right) = \begin{cases} \beta_0 + \mathbf{w}_i\boldsymbol{\lambda} + \mathbf{w}_j\boldsymbol{\lambda}, & \text{if } i \neq j, \\ 0, & \text{if } i = j. \end{cases} \tag{1}$$

This model is equivalent to the site-level component of a GDUM ($v_i, v_j$, see main text), excluding random effects.

##### 2.1.2 Site uniqueness model

Following Legendre and De Cáceres (2013), we define the double-centred matrix as:

$$\mathbf{G} = \mathbf{H}\left(-\frac{1}{2}\mathbf{D}^2\right)\mathbf{H}, \tag{2}$$

where $\mathbf{H} = \mathbf{I} - \frac{1}{n}\mathbf{1}\mathbf{1}^\top$ is the centring matrix, $\mathbf{I}$ is the $n \times n$ identity matrix, $\mathbf{1}$ is an $n$-dimensional vector of ones, and $\mathbf{D}$ is the matrix passed to the double-centring operator. We consider the case where $\mathbf{D}$ is the raw dissimilarity matrix after an element-wise square-root transformation $\mathbf{D} = \sqrt{\mathbf{Y}}$, which is standard for non-Euclidean dissimilarities (Legendre & De Cáceres, 2013). We note that, after substituting $\mathbf{D} = \sqrt{\mathbf{Y}}$, the double centring operator $\mathbf{H}\left(-\frac{1}{2}(\cdot)^2\right)\mathbf{H}$ becomes linear in $\mathbf{Y}$—it can therefore be applied directly to the expected dissimilarity to obtain the expected double-centred matrix:

$$\mathbb{E}(\mathbf{G}) = \mathbf{H}\left(-\frac{1}{2}\mathbb{E}[\mathbf{Y}]\right)\mathbf{H}. \tag{3}$$

The site uniqueness model relates the diagonal elements of $\mathbf{G}$ to environmental predictors:

$$\mathbb{E}\left(\mathrm{SS}_i\right) = \beta_0^* + \mathbf{w}_i\boldsymbol{\lambda}^*, \tag{4}$$

where $\mathrm{SS}_i = g_{ii}$ represents the uniqueness score for site $i$, $\boldsymbol{\lambda}^*$ is the column vector of regression coefficients, and $\beta_0^*$ is the intercept. Note that, for simplicity, we use $\mathrm{SS}_i$ instead of $\mathrm{LCBD}_i = \frac{\mathrm{SS}_i}{\sum_{j=1}^{n}\mathrm{SS}_j}$ as a metric of uniqueness.

#### 2.2 Theoretical derivation

We now show that the uniqueness model coefficients $\boldsymbol{\lambda}^*$ and the dissimilarity model coefficients $\boldsymbol{\lambda}$, are equivalent after a linear transformation.

**Proposition 1.** *Under the pairwise dissimilarity model (Equation 1) and the site* *uniqueness model (Equation 4), the regression coefficients are related by:*

$$\boxed{\boldsymbol{\lambda}^* = \frac{n-2}{n}\boldsymbol{\lambda}}$$

*Proof.* For the diagonal elements of $\mathbf{G}$, the double-centring transformation (Equation 2) yields:

$$\mathrm{SS}_i = g_{ii} = -\frac{1}{2}\left(D_{ii}^2 - \frac{2}{n}\sum_{j=1}^{n} D_{ij}^2 + \frac{1}{n^2}\sum_{j,k=1}^{n} D_{jk}^2\right).$$

Since $D_{ii} = 0$, this simplifies to:

$$\mathrm{SS}_i = \frac{1}{n}\sum_{j=1}^{n} D_{ij}^2 - \frac{1}{2n^2}\sum_{j,k=1}^{n} D_{jk}^2.$$

Substituting the dissimilarity model (Equation 1), we evaluate the required summations to obtain $\mathbb{E}(\mathrm{SS}_i)$. We note that, because the diagonal elements of the expected dissimilarity matrix evaluate to 0 by definition ($D_{ii}^2 = \mathbb{E}(y_{ii}) = 0$) (Equation 1), the case $i = j$ is excluded when replacing $D_{ij}^2$ with the expected dissimilarity $\mathbb{E}(y_{ij})$. The summation is therefore performed over $n-1$ elements for the first summation $\sum_{j=1}^{n} D_{ij}^2$, and over $n(n-1)$ elements for the second summation $\sum_{j,k=1}^{n} D_{jk}^2$.

For the first sum:

$$\begin{aligned}
\sum_{j=1}^{n} D_{ij}^2 &= \sum_{j\neq i} [\beta_0 + \mathbf{w}_i\boldsymbol{\lambda} + \mathbf{w}_j\boldsymbol{\lambda}] \\
&= (n-1)\beta_0 + (n-1)\mathbf{w}_i\boldsymbol{\lambda} + \sum_{j\neq i}\mathbf{w}_j\boldsymbol{\lambda} \\
&= (n-1)\beta_0 + (n-1)\mathbf{w}_i\boldsymbol{\lambda} + (n\overline{\mathbf{w}} - \mathbf{w}_i)\boldsymbol{\lambda} \\
&= (n-1)\beta_0 + (n-2)\mathbf{w}_i\boldsymbol{\lambda} + n\overline{\mathbf{w}}\boldsymbol{\lambda}.
\end{aligned}$$

For the double sum:

$$\sum_{j,k=1}^{n} D_{jk}^2 = \sum_{j \neq k} \left[\beta_0 + (\mathbf{w}_j + \mathbf{w}_k)\boldsymbol{\lambda}\right]$$
$$= n(n-1)\beta_0 + 2(n-1)n\overline{\mathbf{w}}\boldsymbol{\lambda}.$$

Substituting these expressions into the formula for $\mathrm{SS}_i$:

$$\begin{aligned}
\mathbb{E}(\mathrm{SS}_i) &= \frac{1}{n}\left[(n-1)\beta_0 + (n-2)\mathbf{w}_i\boldsymbol{\lambda} + n\overline{\mathbf{w}}\boldsymbol{\lambda}\right] - \frac{1}{2n^2}\left[n(n-1)\beta_0 + 2(n-1)n\overline{\mathbf{w}}\boldsymbol{\lambda}\right] \\
&= \frac{n-1}{n}\beta_0 + \frac{n-2}{n}\mathbf{w}_i\boldsymbol{\lambda} + \overline{\mathbf{w}}\boldsymbol{\lambda} - \frac{n-1}{2n}\beta_0 - \frac{n-1}{n}\overline{\mathbf{w}}\boldsymbol{\lambda} \\
&= \frac{n-1}{2n}\beta_0 + \frac{n-2}{n}\mathbf{w}_i\boldsymbol{\lambda} + \frac{1}{n}\overline{\mathbf{w}}\boldsymbol{\lambda}.
\end{aligned}$$

Rearranging terms yields:

$$\boxed{\mathbb{E}(\mathrm{SS}_i) = \underbrace{\frac{n-1}{2n}\beta_0 + \frac{1}{n}\overline{\mathbf{w}}\boldsymbol{\lambda}}_{\beta_0^*} + \mathbf{w}_i \underbrace{\frac{n-2}{n}\boldsymbol{\lambda}}_{\boldsymbol{\lambda}^*}.} \tag{5}$$

□

#### 2.3 Site-level random effects

In our full hierarchical formulation of GDUM, we include site-level random effects $z_i$, $z_j$. The same linear algebra applies to random effects and fixed effects, which yields

$$\mathbb{E}(\mathrm{SS}_i) = \underbrace{\frac{n-1}{2n}\beta_0 + \frac{1}{n}\overline{\mathbf{w}}\boldsymbol{\lambda}}_{\beta_0^*} + \mathbf{w}_i \underbrace{\frac{n-2}{n}\boldsymbol{\lambda}}_{\boldsymbol{\lambda}^*} + \underbrace{\frac{n-2}{n}z_i}_{\varepsilon_i}. \tag{6}$$

Site-level random effects in the pairwise dissimilarity model can therefore be related to the site-level error $\varepsilon_i$ in the uniqueness model. We note that, if Gaussian distributions are assumed for both site-level random effects and error terms in the site uniqueness model, with $z_i \sim \mathcal{N}(0, \sigma_z^2)$ and $\varepsilon_i \sim \mathcal{N}(0, \sigma_\varepsilon^2)$, the above relation implies that

$$\sigma_\varepsilon^2 = \left(\frac{n-2}{n}\right)^2 \sigma_z^2. \tag{7}$$

### Supplementary figures

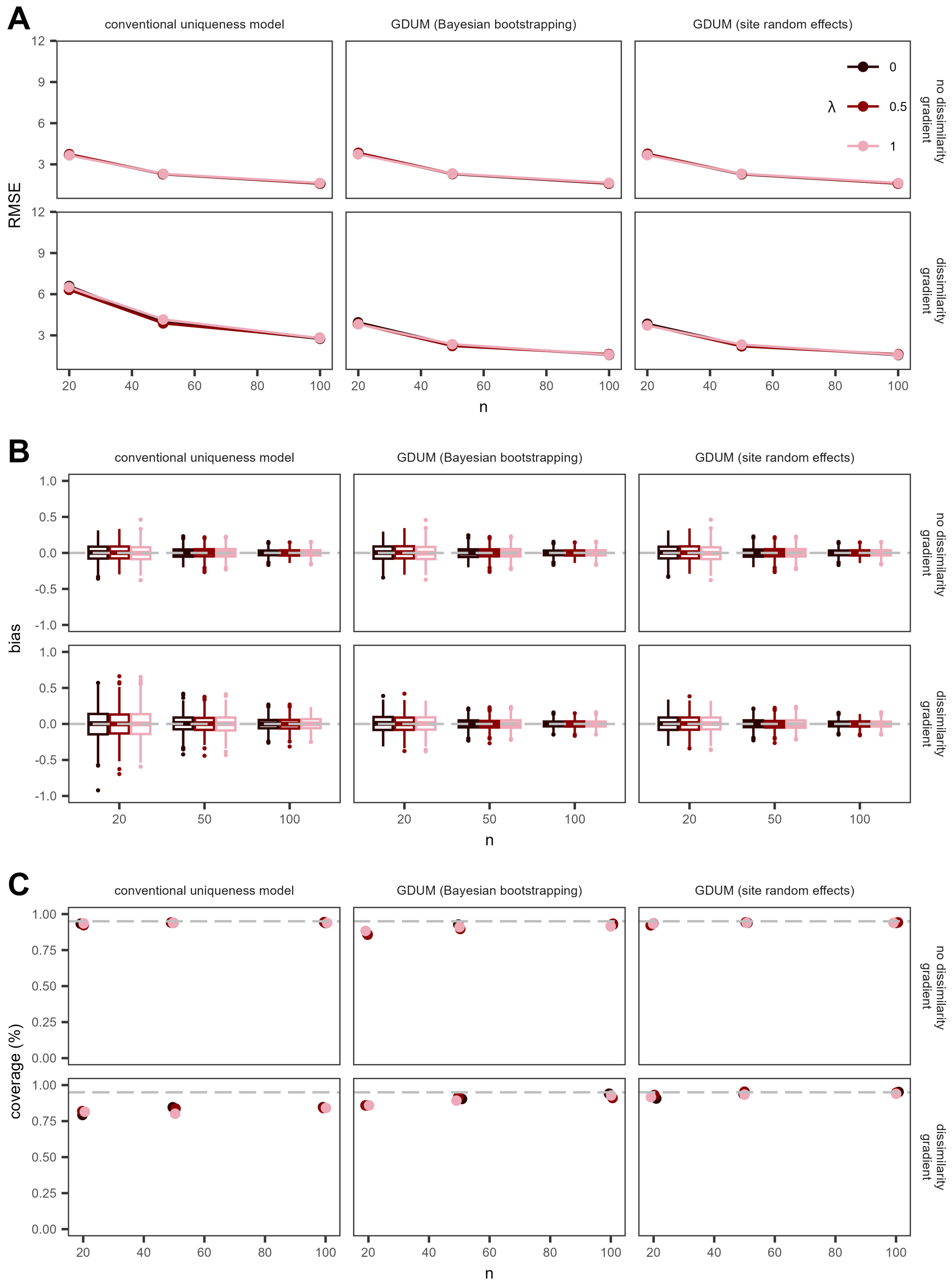


Fig. S2: Performance of GDUMs and conventional uniqueness models with a normally distributed predictor $w$. Different scenarios include varying sample sizes ($n \in \{20, 50, 100\}$), direct effects on uniqueness ($\lambda \in \{0, 0.5, 2\}$), and dissimilarity gradients ($\beta \in \{0, 1\}$). (A) Root Mean Square Error (RMSE) of simulated versus recovered effects on uniqueness ($\lambda$). (B) Bias as the difference between simulated and recovered effects. (C) Parameter coverage as the proportion of simulations where the true, simulated parameter $\lambda$ was within the 95% confidence or credible intervals.

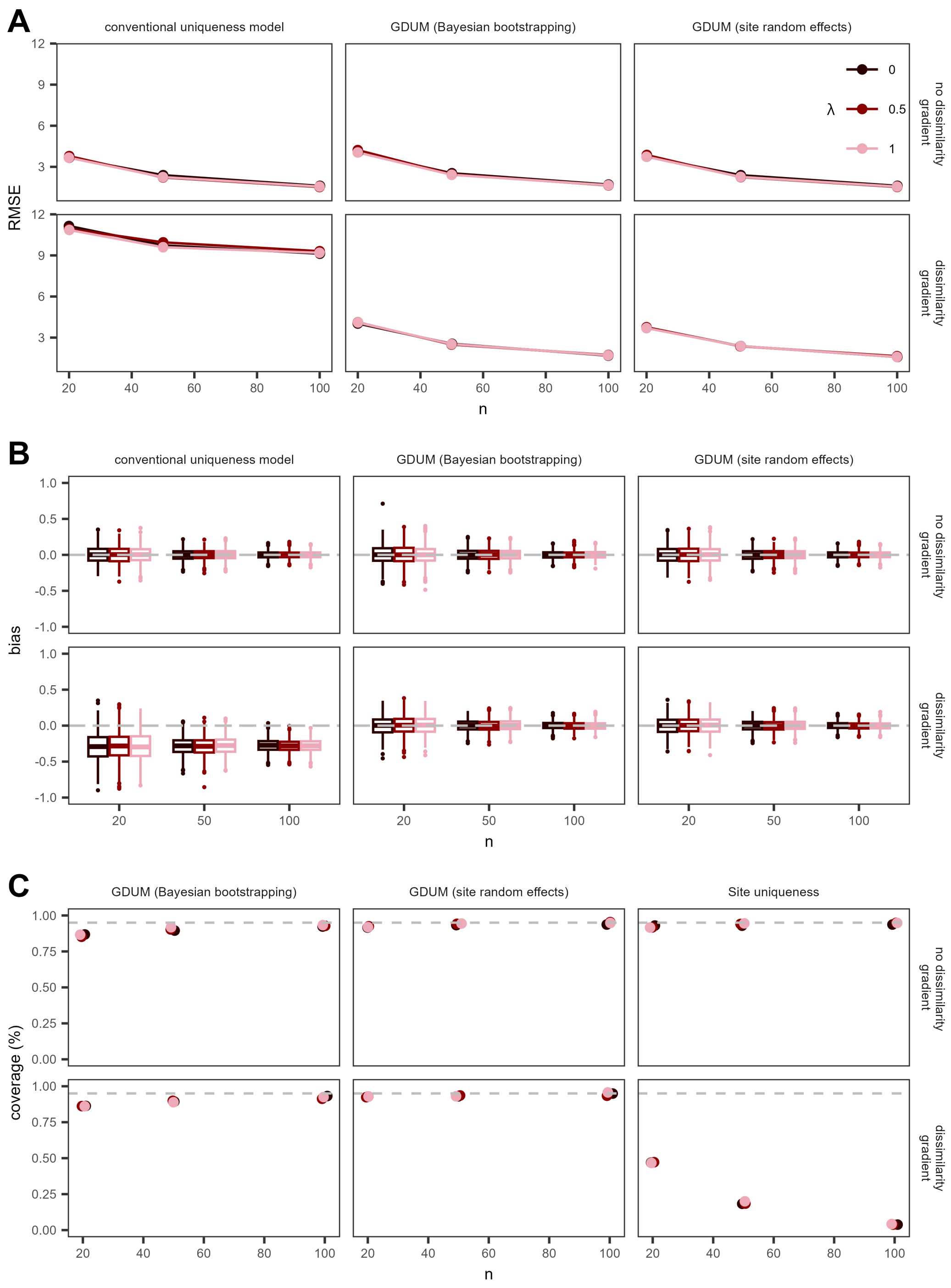


Fig. S3: Performance of GDUMs and conventional uniqueness models with a skewed predictor $w$. Different scenarios include varying sample sizes ($n \in \{20, 50, 100\}$), direct effects on uniqueness ($\lambda \in \{0, 0.5, 2\}$), and dissimilarity gradients ($\beta \in \{0, 1\}$). (A) Root Mean Square Error (RMSE) of simulated versus recovered effects on uniqueness ($\lambda$). (B) Bias as the difference between simulated and recovered effects. (C) Parameter coverage as the proportion of simulations where the true, simulated parameter $\lambda$ was within the 95% confidence or credible intervals.

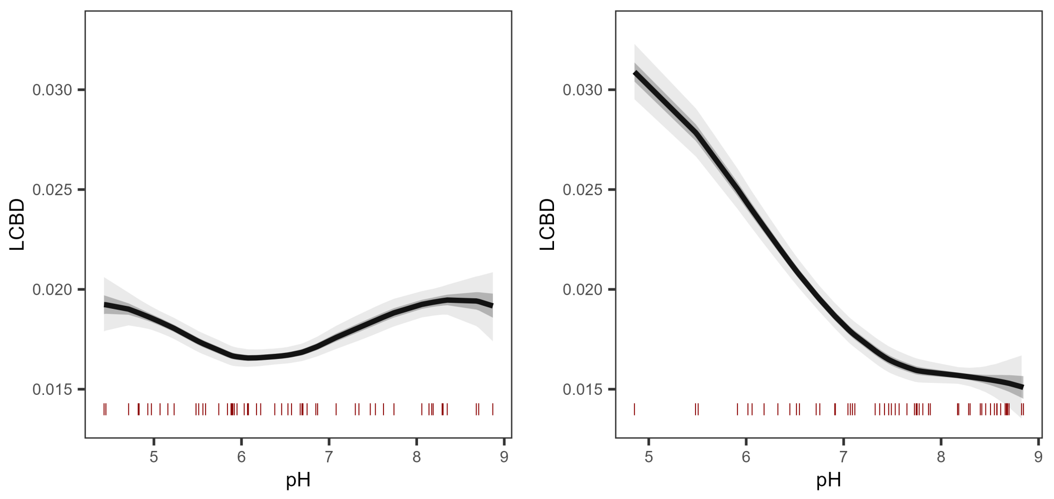


Fig. S4: Contribution of the dissimilarity gradient along pH to the expected LCBD under observed (left) and counterfactual (right) sampling scenarios. LCBD scores were computed from the expected dissimilarity matrix, obtained with the full GDUM model (see section 4.1 in main text). Coloured areas indicate 50% and 95% credible intervals. Red vertical lines indicate the distribution of samples along the pH gradient.

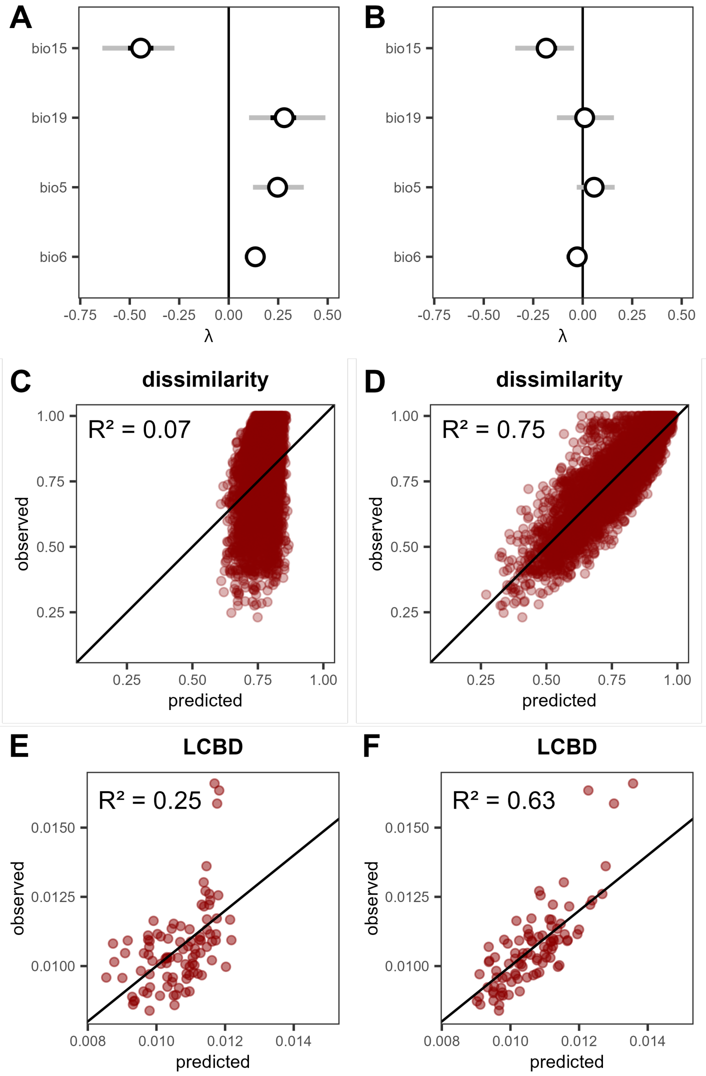


Fig. S5: Comparison of GDUMs with (right panels) and without (left panels) dissimilarity gradients in the pairwise component $h_{ij}$. (A-B) Recovered direct effect of bioclimatic variables on community uniqueness ($\lambda$). Lines represent 50% and 95% credible intervals. (C-D) Model predictions versus observed dissimilarity values. (E-F) LCBD values as derived from model predictions versus computed using the `LCBD.comp` function in the adespatial R package (Dray et al., 2025). Pseudo-$R^2$ based on correlation are displayed indicating goodness of fit.
